# Stress-Enhanced Fear Learning (SEFL) is Associated with Enhanced Reactivation of Fear Engrams in Caudal but not Rostral Dentate Gyrus

**DOI:** 10.64898/2026.03.11.710413

**Authors:** Denisse Paredes, Michael Drew

**Author notes:** **Author e-mails:** Denisse Paredes. **Corresponding Author:**Michael Drew, Ph.D., Center for Learning and Memory University of Texas in Austin Austin, TX, <u>Email:</u> <u>phone</u>.

## Abstract

Traumatic stress can cause long-lasting changes in cognition and affect, sometimes leading to diagnoses such as post-traumatic stress disorder (PTSD). The stress-enhanced fear learning (SEFL) model recapitulates understudied components of PTSD, such as stress-induced sensitization of fear learning. The SEFL procedure entails exposing mice to footshock stress followed later by fear conditioning in a different context. When tested later for recall of fear conditioning, previously stressed mice exhibit enhanced freezing compared to non-stressed controls. Studies have shown that dorsal and ventral dentate gyrus (DG) generates neural ensemble representations of contextual fear, such that fear recall involves reactivation of a sparse set of “engram cells” that were active during fear memory acquisition. How stress affects these hippocampal ensemble representations is unknown. We used SEFL and activity-dependent neuronal tagging with FosTRAP2 mice to investigate effects of stress on fear memory ensembles in rostral and caudal hippocampal DG. FosTRAP2/Ai6 mice received footshock stress or equivalent context exposure without shock in Context A on day 1. Five days later, mice received 1-shock conditioning in Context B and immediately received an injection of 4-OHT (55mg/kg) to tag fear acquisition neurons with the zsGreen reporter. One day later, mice were tested for fear recall in Context B and were perfused 90 minutes after testing. Confirming prior studies, prior stress potentiated 1-shock conditioning in Context B, with stressed mice displaying higher freezing in the Context B test session than non-stressed mice. At the level of neural activity, results showed stress had no effect on the number of zsGreen+ fear ensemble cells or the number of cfos+ recall-activated cells in rostral or caudal DG. However, stress increased reactivation (percentage of zsGreen+ cells expressing cfos) in the caudal but not rostral DG. The results suggest stress potentiates later fear learning by enhancing fear representations in caudal hippocampus, a region of the hippocampus specialized for integrating emotional and motivational valence into memory.

## Introduction

Fear learning is a critical adaptive process that enables organisms to recognize and respond to threats in their environment. However, exposure to trauma can sometimes lead fear responses to become excessive, persistent, or triggered in inappropriate contexts. Individuals with post-traumatic stress disorder (PTSD) often exhibit heightened and generalized fear responses that are resistant to extinction, highlighting the need to understand the underlying neural mechanisms of stress-induced alterations in fear learning (Morey et al., 2015).

The stress enhanced fear learning (SEFL) paradigm is used in rodents to model some of these symptoms of PTSD (Nishimura et al., 2022; Rajbhandari et al., 2018). In the SEFL paradigm, exposure to footshock stress enhances later fear conditioning in a different context (Hassien et al., 2020; Perusini et al., 2016). SEFL is not mediated by generalization of fear between the stress and conditioning contexts, because manipulations that interfere with the stress context memory do not alleviate SEFL. SEFL is thus thought to reflect non-associative sensitization of fear acquisition and/or expression. Although studies indicate that activity and plasticity in the hippocampus is required for the induction of SEFL by stress (Hersman et al., 2019; Pennington et al., 2024), it is currently unknown how stress alters hippocampal neural coding of fear to enhance contextual fear conditioning (CFC).

With the advent of immediate early gene (IEG) trapping techniques to indelibly tag activated neurons within a constrained timeframe, researchers have identified cellular ensembles in the hippocampus that encode fear memories (Denny et al., 2014; Liu et al., 2012; Reijmers et al., 2007). For instance, a sparse ensemble of DG granule cells is activated during acquisition of CFC, and the reactivation of these cells is necessary and sufficient for expression of learned fear (Denny et al., 2014; Liu et al., 2012; Ramirez et al., 2013; Redondo et al., 2014). Optogenetic silencing of the DG fear ensemble decreases expression of learned fear, while optogenetic activation of a DG fear ensemble is sufficient to induce freezing (Liu et al., 2012). Despite these advances, it remains unclear how prior stress exposure modifies the recruitment and reactivation of hippocampal fear memory ensembles, especially along the long axis of the DG (dorsal-rostral to ventral-caudal), which is known to support distinct aspects of memory and emotion (Bannerman et al., 2004; Weeden et al., 2015). In rodents, the dorsal hippocampus is linked to fine-grained spatial cognition and memory (e.g., precise navigation and smaller place fields), whereas the ventral hippocampus is more closely associated with anxiety/emotional behaviors and encodes space at a coarser scale (Bienkowski et al., 2018; Fanselow & Dong, 2010); importantly, evidence supports long-axis gradients rather than a strict dorsal–ventral dichotomy (Ruediger et al., 2012).

Here, we employ the SEFL paradigm and activity-dependent neuronal tagging in FosTRAP2/Ai6 mice to investigate how prior traumatic stress alters the encoding and retrieval of contextual fear memories. We hypothesized that stress would differentially affect the reactivation of fear memory ensembles across the hippocampal long axis. By elucidating the impact of stress on hippocampal memory ensembles, this work aims to advance our understanding of the neural basis for maladaptive fear and inform strategies for treating stress-related disorders.

## Animals

Adult male and female FosTRAP2 (Fostm2.1(icre/ERT2)Luo/J) were crossed with Ai6 (B6.Cg-Gt(ROSA)26Sortm6(CAG-ZsGreen1)Hze/J) mice to generate mice homozygous for FosTRAP2ERT2 allele and homozygous for the zsGreen reporter allele. Breeding pairs for these genotypes were acquired from the Jackson Laboratory (Bar Harbor, ME). Mice were genotyped using polymerase chain reaction (PCR) with protocols from the Jackson Laboratory. A total of 35 mice were used in all experiments (18 females, 17 males, 6-11/group). Mice were weaned at P21 and housed with littermates (3-5 per cage). Mice were housed on a 12-hour light/dark cycle with ad libitum access to food and water. Experiments were performed during the light cycle when mice were 12-15 weeks old. All procedures follow the University of Texas in Austin Institutional Animal Care and Use Committee.

## Neuronal tagging

Recombination was induced as previously described (Zuniga et al., 2024). Briefly, 10mg of 4-hydroxytamoxifen (4-OHT, Sigma Aldrich) was dissolved in a solution of 10% ethanol/90% sunflower seed oil. Immediately after CFC (or context exposure), mice received an intraperitoneal injection of 4-OHT (55mg/kg). Mice were then transported to a satellite housing room with lights off for 72 h. Previous work has demonstrated that housing mice in the dark following 4-OHT injections minimizes non-specific neuronal tagging (Denny et al., 2014). After 72 h, mice returned to the vivarium for the remainder of the experiment.

## Footshock stress session

Mice were handled for 3 days prior to footshock stress. Mice were transported to a holding room adjacent to the behavioral testing room to acclimate for one hour. Mice were then separately moved from their homecage to a conditioning chamber via an opaque container with a clear lid. The stress session consisted of 10 scrambled footshocks (1 mA, 2 secs) over a one-hour session in Context A (narrow grid floors, acetic acid scent on the floor, lights on). Mice then received extinction of Context A for three days (20 minute sessions).

## 1-shock CFC and recall

Three days after the stress session, mice received 1-shock CFC or context exposure. Mice were transported from their home cage to the conditioning chamber via round, plastic containers with a round, clear lid. CFC consisted of one shock (0.75 mA, 2 secs) 3 minutes after the beginning of the session in Context B (wide shock grid, lemon scent, with lights off). Mice were removed from the chamber 30s after the shock. 72 h after conditioning, mice were tested for contextual fear recall in Context B. Freezing was assessed automatically (Video Freeze software, Med Associates) using a linear pixel change algorithm that defined freezing as the absence of movement (Anagnostaras et al., 2010).

## Tissue preparation and immunohistochemistry

90 minutes after contextual fear recall, mice were anesthetized with ketamine and intracardially perfused with 0.1M phosphate buffered saline (PBS) followed by 4% parafolmaldehyde (PFA). Brains were extracted, kept in 4% PFA overnight at 4 C, and transferred to 30% sucrose at 4 C. Tissue was sectioned at 35 um on a cryostat for immunohistochemistry (IHC). About 8-10 coronal sections per mouse including the rostral and caudal hippocampus were used for lHC. The rostral region was collected from -1.22 to -1.94 from Bregma; caudal was-3.16 to -3.80 (Allen Brain Atlas as reference). Sections were washed 3 times in PBS with 0.5% Triton X for 5 minutes. Sections were then blocked at room temperature in 5% normal donkey serum on a shaker. Sections were then incubated overnight at 4 C in primary antibody against c-Fos (1:2000 rat anti c-Fos polyclonal, Synaptic Systems). Sections were next washed in PBS at room temperature (3x, 5 mins) and incubated for an hour in secondary antibody. Next, tissue was washed with PBS at room temperature (3x, 5 mins) and sections were mounted onto slides. As previously described (Zuniga et al., 2024), 3-4 sections per region were imaged at 20x across the z-plane using a Zeiss Axio Imager M2 and counted using ImageJ (Zuniga et al., 2024).

## Results

### Stress increases freezing during contextual fear recall of 1-shock context

Mice received Stress or No Stress (context exposure without footshock) in Context A. Then, 5 days later, mice received 1-shock CFC in Context B or Context B exposure without footshock (No CFC groups). 4-OHT was administered immediately after this session to tag (with zsGreen) cells activated during 1-shock CFC or context exposure. Three days after 1-shock CFC / context exposure, mice received a second exposure to Context B to test for contextual fear expression. Mice in the No CFC groups displayed minimal freezing during both exposures to Context B, so their freezing data is not shown in Figure 1. Prior to footshock in Context B, freezing was low in all groups (Figure 1B “Preshock”), indicating that mice did not generalize fear between the Contexts A and B. An unpaired t-test with Welch’s correction found that stressed and non stressed mice froze at similar levels preshock (t(14)=1.229, p=0.23). Next, we tested whether footshock stress enhances freezing during acquisition and recall of 1-shock CFC. A two tailed t-test also showed that stressed and non-stressed mice froze at similar levels after the shock in the one-shock CFC session (t(14)=0.5649,p=0.5811). In contrast, a two tailed t-test revealed that stressed mice showed increased freezing during the test of CFC recall (t(11.52)=2.475, p=0.03).

**Figure 1.**
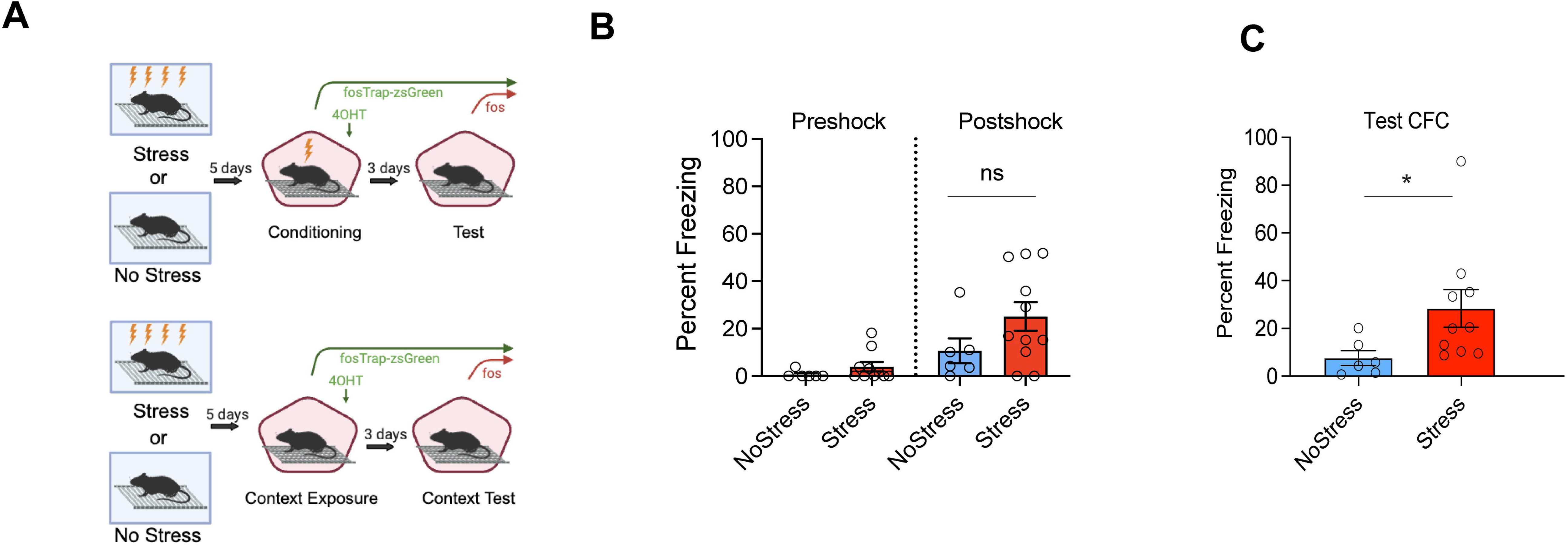
Prior footshock exposure enhances CFC. (A). Behavioral paradigm: mice received either footshock stress or context exposure in context A (left) and then received 1-shock CFC or context exposure alone in context B. Immediately after CFC, all mice received an intraperitoneal injection of 4-OHT. 3 days after conditioning, mice were tested for contextual fear recall in context B. (B) During the CFC session, there were no differences between No-Stress and Stress groups in freezing prior to footshock (t(14)=1.229, p=0.23) (left) and after footshock (t(14)=0.5649, p=0.5811) (middle). (C) During the Contextual fear recall session, Stress mice froze significantly more than No-Stress mice (n=6-10/group; (t(11.52)=2.475, p=0.03).

### Stress does not change the size or reactivation of a CFC ensemble in the rostral dentate gyrus

We investigated whether the number of zsGreen+ neurons (neurons activated during 1-shock CFC or context B exposure) in the rostral and caudal dentate gyrus differed between stressed and non-stressed mice (Figure 2). Four groups were used for these comparisons: 1) No stress-CFC, 2) Stress-CFC, 3) No stress-no CFC and 4) Stress-no CFC. A 2x2 way ANOVA revealed a main effect of CFC on the number of zsGreen+ cells/mm^2^ (F_1,31_=9.565 p=0.0042). The Stress x CFC interaction was not significant (F_1,31_=0.3998, p=0.3998), and there was no main effect of Stress (F_1,31_=3.788 p=0.0607). The same analysis on the number of recall-activated cells (cfos+ cells) showed a significant Stress x CFC interaction (F_1,31_=7.339, p=0.0109), but post-hoc pairwise comparisons between all groups did not detect any significant differences. There were no significant main effects of CFC (F_1,31_=0.2688, p=0.6078) or Stress (F_1,31_=0.3163, p=0.5779). A 2x2 ANOVA was used to assess % reactivation in the rostral DG (percentage of zsGreen+ cells expressing cfos). There was a main effect of CFC (F_1,31_=60.38, p=0.0001), but not a significant CFC x Stress interaction (F_1,31_=2.716, p=0.1095), nor a main effect of Stress (F_1,31_=2.716, p=0.1095). Overall, exposure to CFC (as compared to context exposure without shock) increased the number of zsGreen+ engram cells in DG and increased the reactivation of these zsGreen+ cells during recall. These effects were not modulated by prior stress in rostral DG.

**Figure 2.**
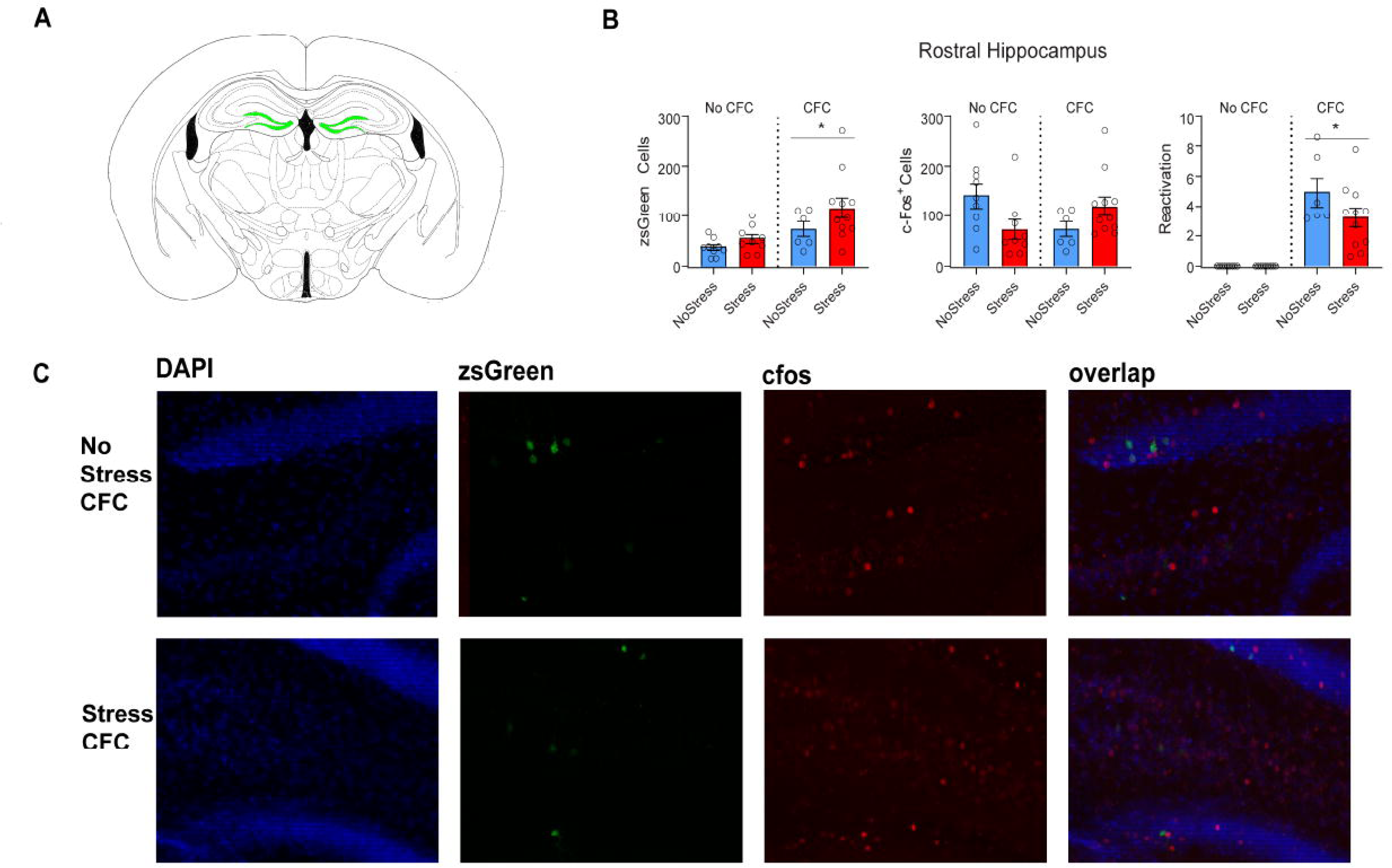
Stress does not change the size or reactivation of a CFC ensemble in the rostral dentate gyrus. (A). Coronal slices displaying (in green) representative locations from which rostral dentate gyrus cell counts were taken. (B). CFC increased the number of zsGreen+ cells (left panel; F_1,31_=9.565 p=0.0042). There was no effect of Stress (F_1,31_=3.788 p=0.0607) or the Stress X CFC interaction (F_1,31_=0.3998, p=0.3998). For recall-activated (cfos+) cells (middle panel) there was a significant Stress x CFC interaction (F_1,31_=7.339, p=0.0109) but not a main effect of CFC (F_1,31_=0.2688, p=0.6078) or Stress (F_1,31_=0.3163, p=0.5779). Similarly, for reactivated cells (percent zsGreen+ cells co-expressing cfos; right panel), there was a main effect of CFC (F_1,31_=60.38, p=0.0001), but no effect of Stress or the Stress X CFC interaction (F_1,31_=2.716, p=0.1095 for both analyses). (C). Representative images showing zsGreen, cfos, and zsGreen/cfos overlap in the rostral DG.

### Stress increases the reactivation of a CFC ensemble in the caudal dentate gyrus

A 2x2 ANOVA on the number of zsGreen+ neurons in the caudal DG (Figure 3) did not reveal a main effect of Stress (F_1,31_=2.809, p=0.1038), nor a main effect of CFC (F_1,31_=0.04591, p=0.8317), nor an interaction (F_1,31_=0.08356, p=0.7744). The same analysis was run on the number of recall-activated cells (cfos+ cells) in the caudal DG, and did not show a significant Stress x CFC interaction (F_1,31_=1.536, p=0.2246), nor a main effect of Stress (F_1,31_=3.451, p=0.0727), nor a main effect of CFC (F_1,31_=1.149, p=0.2920). A 2x2 ANOVA was used to assess % reactivation of tagged zsGreen+ neuronal ensembles in the caudal DG. This analysis revealed a significant interaction (F_1,31_=5.2886, p=0.0284), a main effect of CFC (F_1,31_=36.77, p=0.0001), and a main effect of Stress (F_1,31_=4.546, p=0.0410). Post hoc comparisons revealed a difference between the No Stress-CFC and Stress CFC group (p=0.0198) but not between the two groups that did not receive CFC. Thus, stress increases reactivation of caudal DG fear acquisition neurons but does not modulate activity of a neural ensemble associated with neutral context exposure. Interestingly, correlation analyses between the number of tagged reactivated caudal neurons and contextual recall freezing was not significant (r(8)=-0.04728, p=0.8968; data not shown).

**Figure 3.**
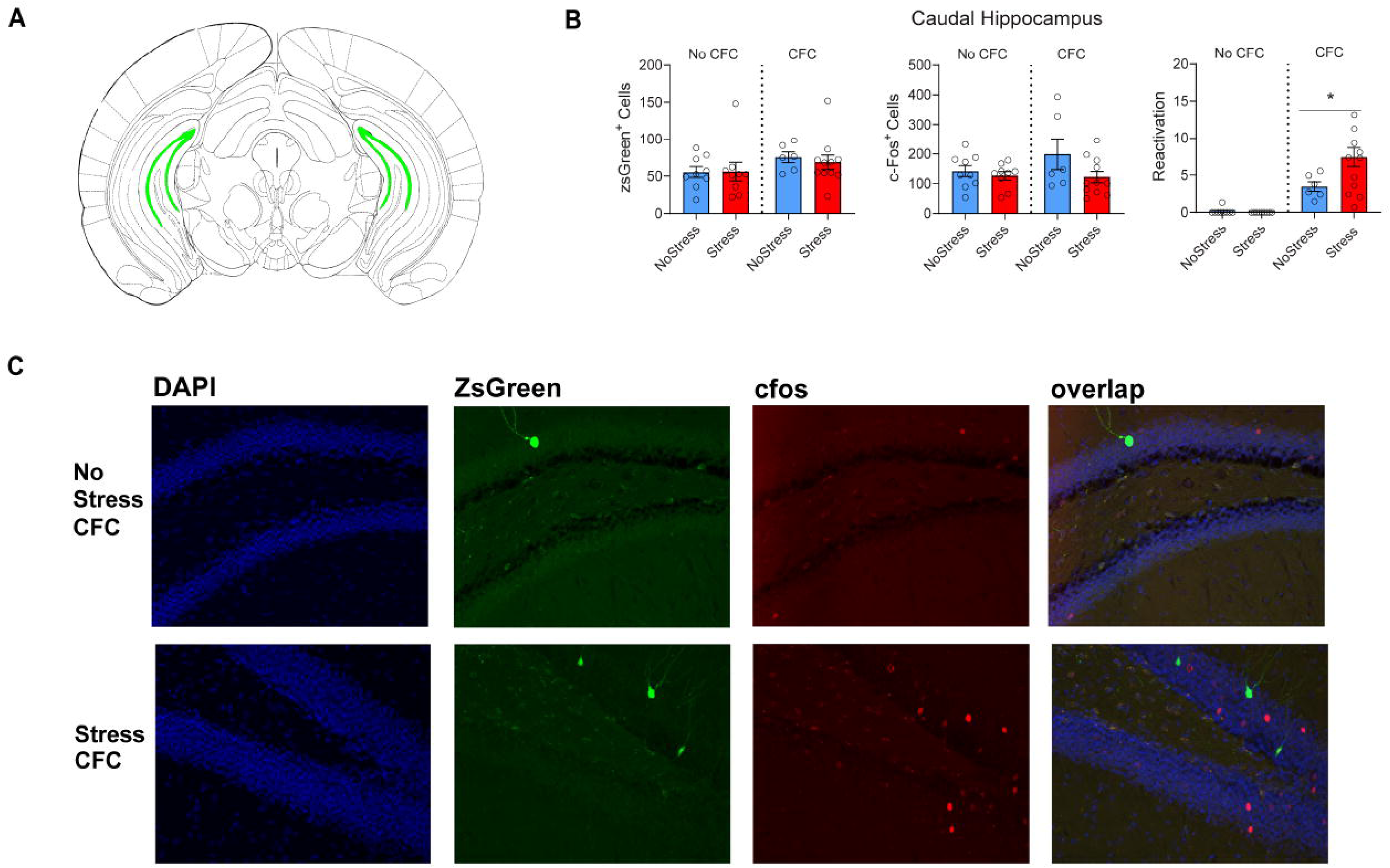
Prior stress increases the reactivation of a CFC ensemble in the caudal dentate gyrus. (A). Coronal slices displaying (in green) representative locations from which caudal dentate gyrus cell counts were taken. (B). There was no effect of Stress (F_1,31_=2.809, p=0.1038), CFC (F_1,31_=0.04591, p=0.8317), or the Stress X CFC interaction on the number of zsGreen+ neurons in the caudal DG (F_1,31_=0.08356, p=0.7744) (left panel). There was no effect of Stress (F_1,31_=3.451, p=0.0727), nor a main effect of CFC (F_1,31_=1.149, p=0.2920), nor a Stress x CFC interaction effect (F_1,31_=0.08356, p=0.7744) on the number of cfos+ cells (middle panel). For reactivated cells, there was a main effect of Stress (F_1,31_=4.546, p=0.0410), a main effect of CFC (F_1,31_=36.77, p=0.0001), and a Stress x CFC interaction (F_1,31_=5.2886, p=0.0284). Stress enhances % reactivation during recall in the caudal DG in fear conditioned animals (right, p=0.0198). (C). Representative images showing zsGreen, cfos, and zsGreen/cfos overlap in the caudal DG.

## Discussion

Our results confirm that prior exposure to stress enhances CFC in a novel context. Additionally, using activity-dependent neuronal tagging, we investigated how stress affects the establishment and reactivation of neural ensemble representations of contextual fear in the rostral and caudal DG. The main findings were that prior stress had no effect on the number of rostral DG or caudal DG neurons activated during acquisition of CFC, but, during CFC recall, stressed mice displayed enhanced reactivation of fear acquisition neurons in caudal DG but not rostral DG. The enhancement of ensemble reactivation was specific to a CFC memory; stress did not enhance reactivation of an ensemble coding a neutral (unshocked) context exposure. Because reactivation of fear acquisition ensembles in the DG is necessary and sufficient for expression of CFC (Denny et al., 2014; Lacagnina et al., 2019; Liu et al., 2012)the increased reactivation in caudal DG may provide a mechanism through which stress enhances learned fear.

These results suggest that stress enhances caudal hippocampal coding of contextual fear but does not modulate rostral hippocampal coding. In our study, rostral hippocampus included what is traditionally defined as dorsal hippocampus, whereas caudal was mainly ventral DG (although it also included portions of dorsal DG). Our findings thus comport with previous work demonstrating that although both dorsal and ventral hippocampus contribute to contextual fear learning (Huckleberry et al., 2018), these regions appear to play different roles. The dorsal hippocampus is crucial for spatial representations of context (Matus-Amat et al., 2004) while the ventral region modulates innate anxiety and defensive behaviors (Turner et al., 2022). The dorsal hippocampus generates highly specific and detailed representations of spatial information, which are critical for distinguishing between similar environments or experiences. Compared to neurons in the dorsal region, ventral hippocampal neurons have larger place fields and carry less spatial information (Jung et al., 1994). Further, ventral hippocampus interacts extensively with limbic structures like the amygdala and hypothalamus, enabling it to integrate affective information into memory (Ciocchi et al., 2015; Gergues et al., 2020). Our findings suggest that the contextual fear representations in the caudal, but not rostral DG may underlie the SEFL phenotype. Increased reactivation of hippocampal fear memory ensembles enhances fear memory generalization to different contexts (Bian et al., 2019; Cui et al., 2024). Thus, it is possible that stress-induced enhancement of fear ensemble reactivation may increase fear generalization. This hypothesis requires testing.

We also observed an interaction between stress and fear conditioning in rostral DG, such that stress trended toward having opposite effects on cFos expression in mice that were and were not fear conditioned. It is difficult to interpret these data since the pairwise comparisons did not reach significance. Previous studies show that stress can have bidirectional effects on hippocampal IEG expression. For instance, chronic restraint elevates basal c-FOSand ArcmRNAs 24 hours after the final session(Pacheco et al., 2017). However, other work indicates that repeated stress can blunt or habituate IEG responses(Ostrander et al., 2009). Our results raise the possibility that stress effects on hippocampal IEG expression could be modulated by subsequent fear conditioning, but further studies will be needed to confirm this.

Our findings are congruent with other work describing differing effects of stress on dorsal and ventral hippocampus. In particular, stress can modulate hippocampal plasticity in opposite patterns along the dorsal and ventral axis. Stress, suppresses long-term potentiation in the dorsal pole, while enhancing potentiation in the ventral pole (Maggio & Segal, 2007), lending credence to the idea that stress could enhance plasticity in vDG during fear conditioning leading to greater reactivation of fear ensembles in caudal hippocampus. The dorsal and ventral hippocampus also exhibit differing molecular responses to acute stress, (von Ziegler et al., 2022). The ventral hippocampus integrates information from several stress sensitive regions such as the BLA (Turner et al., 2022). The BLA has been identified as a key region mediating SEFL, such that BLA inhibition during stress ablates enhanced fear recall (Perusini et al., 2016). BLA input to the ventral hippocampus also modulates fear memory consolidation, and activation of BLA-ventral hippocampus promotes anxiety-like behaviors (Felix-Ortiz et al., 2013; Huff et al., 2016). Therefore, it is possible that BLA input to the ventral hippocampus is necessary for stress enhancement of fear retrieval. The ventral hippocampus may also modulate stress-enhanced fear through its outputs, such as the amygdala and prefrontal cortex (Jin & Maren, 2015). The prefrontal cortex is a stress sensitive region that modulates fear conditioning and extinction (Giustino & Maren, 2015). Fear memory ensembles have been identified in the prefrontal cortex (Cummings & Clem, 2020; Zelikowsky et al., 2014). Stimulation of the prelimbic region of the prefrontal cortex enhances fear expression (Vidal-Gonzalez et al., 2006). Therefore, future work should focus on stress effects on reactivation of fear ensembles in other regions mediating associative fear learning, such as the amygdala and prefrontal cortex.

This work uncovers a neural ensemble in the caudal dentate gyrus that may contribute to the stress enhanced fear learning phenotype in rodents. We demonstrate that prior stress sensitizes the expression of new learned fear, and that this effect is associated with increased reactivation of a putative engram in the caudal dentate gyrus. Importantly, we have identified that stress induces differing effects in the reactivation of contextual fear memory reactivation in the caudal versus rostral dentate gyrus. Our results suggest stress may enhance fear learning expression via a caudal hippocampal-dependent mechanism. Given the distinct circuitry between the rostral and caudal hippocampus, stress may influence the caudal hippocampus via interactions with other stress-sensitive regions, such as the prefrontal cortex and BLA. As discussed previously, the increased reactivation of fear engram cells in the caudal dentate gyrus may reflect the propensity of stress to induce generalization of the encoded fear memory. Our work provides rationale to explore a caudal-hippocampal dependent mechanism of stress-induced fear generalization.

## Acknowledgements

We thank members of the Drew lab for support and input. This work was funded by R01 MH117426.

